# Unified Multi-Cohort Harmonisation and Normative Modelling of Neuroimaging Data via Hierarchical GAMLSS

**DOI:** 10.64898/2026.03.08.710422

**Authors:** Mai P. Ho, Nikita K. Husein, Lei Fan, Rachel Visontay, Hollie Byrne, Emma K. Devine, Lindsay M. Squeglia, Perminder S. Sachdev, Jiyang Jiang, Wei Wen, Louise Mewton

## Abstract

Large-scale neuroimaging studies increasingly pool data across multiple cohorts, scanners, and acquisition protocols, introducing technical between-cohort variation that must be addressed before meaningful biological inference can be drawn. Existing harmonisation methods, particularly ComBat-based approaches, have been widely adopted for this purpose. However, they remain limited by Gaussian assumptions and by their focus on location or location-scale correction. In this study, we propose a unified hierarchical Generalised Additive Models for Location, Scale and Shape (GAMLSS) framework for multi-cohort harmonisation and normative modelling of structural neuroimaging data. The framework models cohort effects directly within all fitted distributional parameters, accommodates any parametric family for which exact inverse mapping is available, and returns harmonised values on the original measurement scale through centile-based quantile mapping. Normative deviation scores are obtained as a direct by-product of the same fitted model, enabling harmonisation and normative inference to be conducted jointly. The method was evaluated in a pooled longitudinal dataset comprising 88,126 observations across 237 structural neuroimaging features from six cohorts spanning childhood to late life: ABCD, IMAGEN, NCANDA, LIFE, UK Biobank, and MAS. Harmonisation performance was compared with ComBat, ComBat-GAM, and ComBat-LS using complementary criteria assessing data retention, residual batch effects, preservation of age-related and sex-related biological signal, and coherence of post-harmonisation lifespan trajectories. GAMLSS achieved near-complete removal of residual cohort effects, retained almost all valid observations post-harmonisation, and showed the strongest overall preservation of biological signal across validation metrics. In particular, it better preserved biologically plausible age trajectories for distributionally complex features such as white matter hypointensity volume, while simultaneously providing harmonised native-scale values and normative deviation scores within a single framework. These findings suggest that hierarchical GAMLSS offers a flexible and practical alternative to existing ComBat-based methods for large-scale neuroimaging harmonisation, particularly for features with non-Gaussian residual distributions and settings where cohort effects extend beyond differences in mean and variance.

## 1. Introduction

Large multi-site neuroimaging studies are increasingly used to improve statistical power, broaden age coverage, and support more generalisable models of brain structure across the lifespan. However, pooling MRI data across cohorts, scanners, and acquisition protocols introduces unwanted technical variation that can distort biological inference if left unaddressed. In structural neuroimaging, these batch effects can arise from differences in scanner hardware, acquisition parameters, image processing pipelines, and other study-specific factors, and they may obscure or bias associations with age, sex, and disease-related variation (Han et al., 2006; Jovicich et al., 2006; Takao et al., 2014). Harmonisation therefore aims to reduce non-biological between-study variability while preserving meaningful biological signal (Fortin et al., 2017; Johnson et al., 2007).

The dominant harmonisation framework in neuroimaging has been ComBat. Originally developed for genomics, ComBat uses an empirical Bayes approach to estimate and remove additive batch effects in the mean and multiplicative batch effects in the variance while retaining specified biological covariates (Johnson et al., 2007). Fortin and colleagues subsequently adapted ComBat to neuroimaging, helping establish it as a standard method for retrospective harmonisation in pooled imaging studies (Fortin et al., 2017, 2018). In its classical form, however, ComBat is still fundamentally a location-scale correction model. It assumes a parametric form for the batch effect and is typically paired with linear covariate adjustment. As a result, a number of ComBat variants have been developed to address specific limitations of the classical model. ComBat-GAM extends the framework to accommodate non-linear age effects through generalised additive modelling, which is essential for lifespan neuroimaging where many brain measures follow non-linear developmental and ageing trajectories (Pomponio et al., 2020). Longitudinal ComBat (ComBat-Long) incorporates subject-level random intercepts to account for repeated measurements and within-person correlation in longitudinal data (Beer et al., 2020). Other extensions include CovBat for covariance harmonisation in multivariate settings and ComBat-seq for count data (A. A. Chen et al., 2022; Zhang et al., 2020). Despite these developments, earlier ComBat variants still harmonise scale effects by rescaling variance towards a common site-adjusted distribution, rather than explicitly preserving covariate-dependent heteroskedasticity in the adjusted data. More recently, ComBat-LS was proposed to address this limitation by preserving biological variation in both feature location and scale (Gardner et al., 2025). These developments have substantially broadened the ComBat family. However, most existing harmonisation methods still remain centred on correcting batch effects in location and scale. This may be restrictive for structural neuroimaging phenotypes, whose conditional distributions often depart from Gaussian assumptions through heteroskedasticity, skewness, and kurtosis (de Boer et al., 2024). Consequently, site or cohort effects may also extend beyond differences in mean and variance alone. This suggests a need for more flexible harmonisation frameworks that can model the full conditional distribution, including higher-order moments where supported by the data.

A related but conceptually distinct line of work comes from normative modelling. Rather than treating site variation purely as a preprocessing problem, normative models estimate a covariate-conditional reference distribution and quantify how far each individual deviates from the expected range, typically using centiles or z-scores (Rutherford et al., 2022). A notable example is the hierarchical Bayesian regression normative framework, implemented in PCNtoolkit, which accommodates site variation directly within the normative model rather than through a separate harmonisation step (Bayer et al., 2022; de Boer et al., 2024; Kia et al., 2020). This approach is motivated by the concern that site is often entangled with biological variables of interest, so two-stage correction can remove meaningful signal, distort predictive variance, and bias downstream inference. In particular, hierarchical Bayesian regression (HBR) models site as a random effect with partial pooling across sites, allowing site-specific parameters to borrow strength from a shared prior. Bayer et al. reported that this single-stage strategy produced better-calibrated normative predictions and z-scores with little to no residual site effect, while two-stage harmonisation models showed substantial variance shrinkage (Bayer et al., 2022). However, the primary output of these frameworks is usually a site-adjusted centile or deviation score rather than a harmonised measurement returned on the original feature scale. This can be limiting in applications that still require native-scale data, including descriptive visualisation, standard regression analyses, and downstream pipelines that operate on the original units.

In this study, we introduce a unified hierarchical Generalised Additive Models for Location, Scale and Shape (GAMLSS) framework for multi-site harmonisation and normative modelling of structural neuroimaging data. Extending the generalised additive models (GAM), GAMLSS are a flexible class of regression models in which multiple parameters of the outcome distribution can be modelled as functions of covariates, allowing the conditional distribution to vary in location, scale, and shape (Rigby & Stasinopoulos, 2005). This makes GAMLSS well suited to neuroimaging applications where outcomes may show non-linearity, heteroskedasticity, skewness, and kurtosis, and has led to its use in normative modelling studies including large-scale lifespan brain charts (Bethlehem et al., 2022; Zhuo et al., 2025). Building on this foundation, our framework models cohort effects directly within the relevant distributional parameters, allowing adjustment of location, scale, and higher-order moments of the conditional distribution. Harmonised values are obtained by removing cohort-specific random effects and mapping observations back to the original measurement scale through centile-based quantile mapping, while the same fitted model is used to derive normative deviation scores.

We applied this framework to pooled longitudinal structural MRI data from six cohorts spanning childhood to late life, including the Adolescent Brain Cognitive Development Study (ABCD) (Volkow et al., 2018), IMAGEN (Schumann et al., 2010), the National Consortium on Alcohol and Neurodevelopment in Adolescence (NCANDA) (Brown et al., 2015), the LIFE Adult Study (LIFE) (Loeffler et al., 2015), the UK Biobank (Sudlow et al., 2015), and the Sydney Memory and Ageing Study (MAS) (Sachdev et al., 2010). The combined dataset comprises 64,086 baseline participants and 88,126 observations in total. The analysis includes 237 structural neuroimaging features grouped into four categories: cortical volumes, cortical thickness, cortical surface area, and subcortical/global volumes derived from FreeSurfer automated segmentation. We benchmark the proposed framework against classical ComBat, ComBat-GAM, and ComBat-LS using a two-component validation strategy that evaluates both batch-effect removal and the preservation of biological signals. In doing so, this study tests whether a single flexible distributional framework can jointly provide harmonised original-scale data and normative deviation scores. It also addresses limitations of existing approaches, which either focus primarily on location-scale correction or produce deviation scores without harmonised outputs on the native measurement scale.

## 2. Materials and methods

### 2.1 Study population and cohort description

Data were drawn from six independent neuroimaging studies spanning the human lifespan, each with a prospective longitudinal design, yielding a combined analytic sample of N = 64,086 participants (age range: 8.9–95.9 years, 88,126 observations in total). These cohorts were selected to provide broad and overlapping age coverage across childhood, adolescence, adulthood, and late life. All six studies contributed T1-weighted structural MRI data acquired on 3T scanners alongside demographic information including age, sex, and scanner identifiers. Analytic sample sizes reflect participants retained after quality control and data availability checks and may therefore differ from total cohort recruitment figures. A summary of baseline characteristics is presented in Table 1, and the longitudinal data structure across cohorts is described in Table 2.

**Table 1.** Baseline participant characteristics (N participants = 64,086)

|  | **Total** | **ABCD** | **IMAGEN** | **NCANDA** | **LIFE** | **UKB** | **MAS** |
| --- | --- | --- | --- | --- | --- | --- | --- |
| **Sample size** |  |  |  |  |  |  |  |
| Participants, n | 64,086 | 11,267 | 2,090 | 806 | 2,537 | 46,850 | 536 |
| **Demographics** |  |  |  |  |  |  |  |
| Male, number (%) | 30,979 (48.3) | 5,841 (51.8) | 1,027 (49.1) | 395 (49.0) | 1,340 (52.8) | 22,134 (47.2) | 242 (45.1) |
| Age at baseline, years, mean ±SD | 52.8 ± 23.5 | 9.9 ± 0.6 | 14.4 ± 0.4 | 16.2 ± 2.5 | 59.2 ± 15.0 | 64.7 ± 7.7 | 78.6 ± 4.7 |
| Age at baseline, years, range [min, max] | [8.9, 90.6] | [8.9, 11.1] | [10.8, 16.0] | [12.0, 22.0] | [19.1, 83.0] | [44.6, 83.7] | [70.4, 90.6] |
| **Design/Acquisition** |  |  |  |  |  |  |  |
| Within-cohort batch, n (levels) | 49 | 29 | 8 | 5 | 1 | 4 | 2 |
| Participants per batch, n, median [IQR] | 332 [208–503] | 421 [229–540] | 258 [255–261] | 157 [148–171] | 2,537 [2,537–2,537] | 9650 [5,371–15,991] | 268 [263–273] |

SD = standard deviation; IQR = interquartile range. Baseline is the first available visit per participant and does not require wave labels to align across cohorts. Detailed batch composition and counts are reported in Supplementary 2.

**Table 2.**
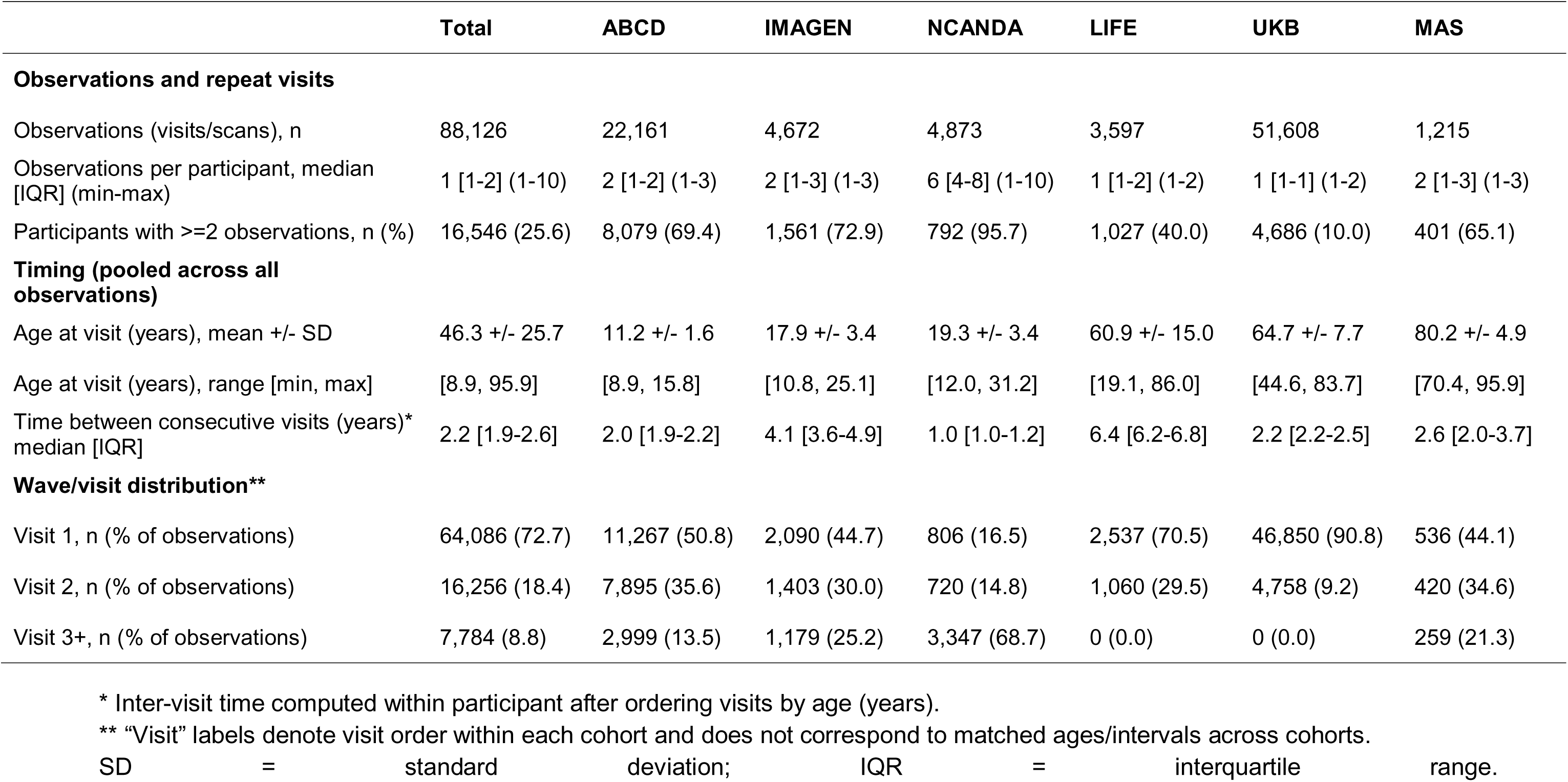
Longitudinal data structure and follow-up (all available observations)

|  | Total | ABCD | IMAGEN | NCANDA | LIFE | UKB | MAS |
| --- | --- | --- | --- | --- | --- | --- | --- |
| **Observations and repeat visits** |  |  |  |  |  |  |  |
| Observations (visits/scans), n | 88,126 | 22,161 | 4,672 | 4,873 | 3,597 | 51,608 | 1,215 |
| Observations per participant, median [IQR] (min-max) | 1 [1-2] (1-10) | 2 [1-2] (1-3) | 2 [1-3] (1-3) | 6 [4-8] (1-10) | 1 [1-2] (1-2) | 1 [1-1] (1-2) | 2 [1-3] (1-3) |
| Participants with >=2 observations, n (%) | 16,546 (25.6) | 8,079 (69.4) | 1,561 (72.9) | 792 (95.7) | 1,027 (40.0) | 4,686 (10.0) | 401 (65.1) |
| **Timing (pooled across all observations)** |  |  |  |  |  |  |  |
| Age at visit (years), mean +/- SD | 46.3 +/- 25.7 | 11.2 +/- 1.6 | 17.9 +/- 3.4 | 19.3 +/- 3.4 | 60.9 +/- 15.0 | 64.7 +/- 7.7 | 80.2 +/- 4.9 |
| Age at visit (years), range [min, max] | [8.9, 95.9] | [8.9, 15.8] | [10.8, 25.1] | [12.0, 31.2] | [19.1, 86.0] | [44.6, 83.7] | [70.4, 95.9] |
| Time between consecutive visits (years)* median [IQR] | 2.2 [1.9-2.6] | 2.0 [1.9-2.2] | 4.1 [3.6-4.9] | 1.0 [1.0-1.2] | 6.4 [6.2-6.8] | 2.2 [2.2-2.5] | 2.6 [2.0-3.7] |
| **Wave/visit distribution\*\*** |  |  |  |  |  |  |  |
| Visit 1, n (% of observations) | 64,086 (72.7) | 11,267 (50.8) | 2,090 (44.7) | 806 (16.5) | 2,537 (70.5) | 46,850 (90.8) | 536 (44.1) |
| Visit 2, n (% of observations) | 16,256 (18.4) | 7,895 (35.6) | 1,403 (30.0) | 720 (14.8) | 1,060 (29.5) | 4,758 (9.2) | 420 (34.6) |
| Visit 3+, n (% of observations) | 7,784 (8.8) | 2,999 (13.5) | 1,179 (25.2) | 3,347 (68.7) | 0 (0.0) | 0 (0.0) | 259 (21.3) |

\* Inter-visit time computed within participant after ordering visits by age (years).
\*\* "Visit" labels denote visit order within each cohort and does not correspond to matched ages/intervals across cohorts.
SD = standard deviation; IQR = interquartile range.

#### Adolescent Brain Cognitive Development Study (ABCD)

The ABCD Study is the largest long-term study of brain development and child health in the United States (Casey et al., 2018; Volkow et al., 2018). Between 2016 and 2018, a multi-site consortium of 21 data acquisition sites recruited approximately 11,880 children aged 9–10 years using a stratified probability sample of schools, with the aim of following participants through adolescence into young adulthood. Comprehensive assessments span brain imaging, neurocognition, physical and mental health, substance use, and cultural and environmental factors. The imaging protocol was harmonised across three 3T scanner platforms (Siemens Prisma, GE MR750, and Philips); multiband echo-planar imaging sequences were used for functional and diffusion acquisitions, and a high-resolution T1-weighted magnetisation-prepared rapid gradient echo sequence was acquired for structural imaging (Casey et al., 2018; Hagler et al., 2019). In the present analysis, we included n = 11,267 baseline participants across up to three longitudinal waves, yielding 22,161 total observations. Within-cohort batch effects were defined by unique combinations of acquisition site and scanner configuration, resulting in 32 batches.

#### IMAGEN

IMAGEN is a European multi-centre longitudinal imaging genetics study designed to investigate the genetic and neurobiological basis of individual differences in reinforcement sensitivity, impulsivity, and emotional reactivity during adolescence (Mascarell Maričić et al., 2020; Schumann et al., 2010). Approximately 2,000 adolescents were recruited at age 14 from schools across eight sites in the United Kingdom, Ireland, France, and Germany, with follow-up assessments at ages 16, 19, and 22. We included n = 2,090 baseline participants across up to three waves, yielding 4,672 total observations across 8 recruiting sites.

#### National Consortium on Alcohol and Neurodevelopment in Adolescence (NCANDA)

NCANDA is a US-based accelerated longitudinal study designed to disentangle the relationships between alcohol use and neurodevelopmental change during adolescence and young adulthood (Brown et al., 2015). A sample of 831 youth aged 12–21 years was recruited across five sites, with the majority (83%) having minimal or no prior alcohol exposure at baseline. Data were acquired on 3T systems from two manufacturers (GE Discovery MR750 and Siemens TIM Trio). We included n = 806 baseline participants across up to 10 longitudinal waves, yielding 4,873 total observations. This is the densest longitudinal sampling of any cohort in this study, with 95.7% of participants contributing two or more observations. Within-cohort batch effects were defined by unique combinations of site and scanner configuration, yielding 10 batches.

#### LIFE-Adult-Study (LIFE)

The LIFE-Adult-Study is a population-based cohort conducted by the Leipzig Research Centre for Civilization Diseases, University of Leipzig, Germany (Engel et al., 2023; Loeffler et al., 2015). Adults primarily aged 40–79 years were recruited from the city of Leipzig to investigate lifestyle and environmental determinants of major civilisation diseases including cardiovascular, metabolic, and psychiatric disorders. Brain imaging was performed on a 3T Siemens Verio scanner. We included n = 2,537 baseline participants across up to two assessment waves, yielding 3,597 total observations.

#### UK Biobank (UKB)

UK Biobank is a large-scale population-based prospective study that recruited 503,317 participants aged 40–69 years between 2006 and 2010 across the United Kingdom (Sudlow et al., 2015). Beginning in 2014, a subset of participants was re-invited for multi-modal imaging, including brain, cardiac, and abdominal MRI, as part of the world’s largest imaging study, with a proportion of imaged participants also undergoing a repeat imaging visit (Littlejohns et al., 2020; Miller et al., 2016). Brain imaging is performed on dedicated 3T Siemens Skyra scanners at purpose-built imaging centres using a standardised and highly controlled acquisition protocol. In the present analysis, we included n = 51,608 participants (47.3% male; age range: 44.6–83.7 years) across up to two imaging visits acquired on 4 sites.

#### Sydney Memory and Ageing Study (MAS)

The Sydney MAS is an epidemiological cohort study of brain ageing and dementia initiated in 2005 (Sachdev et al., 2010). Non-demented community-dwelling adults aged 70–90 years were recruited from eastern Sydney via random approach to the electoral roll and underwent biennial follow-up assessments including structural MRI acquired on Philips 3T systems. We included n = 536 baseline participants across up to three waves acquired on 2 scanners, yielding 1,215 total observations.

### 2.2 Feature derivation and data pre-processing

Structural T1-weighted MRI data from all cohorts were processed using FreeSurfer (Fischl, 2012), applying the Desikan-Killiany (DK) atlas (Desikan et al., 2006) for cortical parcellation and the automated subcortical segmentation (ASEG) atlas for subcortical structures. A total of 237 structural neuroimaging features were extracted across four categories: subcortical and global volumes (n = 37), capturing volumetric measures in mm³ of subcortical structures, ventricles, and global brain compartments including white matter hypointensities; cortical volumes (n = 66) and cortical thickness (n = 68), capturing regional volumes in mm³ and mean thickness in mm across 34 bilateral DK atlas parcels; and cortical surface area (n = 66), capturing regional surface area in mm² across the same parcels. Features with negative values or missing data for age, sex, cohort, or within-cohort batch were excluded prior to harmonisation. A full feature list with abbreviations and descriptions is provided in Supplementary Table 1.

To reduce scanner-related technical variation within cohorts while accounting for repeated measurements, we first applied ComBat-Long separately within each cohort. This approach was used because a single model that simultaneously captured within-cohort scanner effects, between-cohort differences, and subject-level random effects was not computationally feasible given the complexity and heterogeneity of the longitudinal designs across cohorts. Applying ComBat-Long within cohort therefore addressed scanner-related variation at the site level while preserving the higher-level cohort differences targeted in the subsequent harmonisation step. This distinction is important because within-cohort scanner effects reflect technical variation between acquisition sites, whereas between-cohort differences may also capture broader sources of heterogeneity, including recruitment, protocol, and population differences.

### 2.3 Empirical motivation analysis

To empirically motivate the need for flexible distributional modelling in cross-cohort harmonisation, we examined between-cohort distributional differences using baseline observations from UKB and LIFE within their overlapping age range of 44 to 83 years, yielding 46,842 baseline UKB participants and 2,063 baseline LIFE participants. This analysis was conducted on post-ComBat-Long data to isolate genuine cross-cohort batch effects from within-cohort scanner artefacts.

Three complementary analyses were conducted. First, a batch effect R-squared was computed for each feature to quantify the magnitude of cohort-related variance. Two linear models were fitted: a baseline model containing age and sex, and an extended model additionally including cohort. The increment in R-squared when cohort is added reflects the additional variance attributable to the cohort batch effect independent of biological covariates. Second, between-cohort differences in distributional moments were quantified for each feature using three measures: the absolute log standard deviation (SD) ratio to capture differences in scale, the absolute delta skewness to capture differences in asymmetry, and the absolute delta excess kurtosis to capture differences in tail heaviness. Log SD ratio was used in place of the raw SD difference because it is scale-invariant across features with different measurement units. All three measures reflect the magnitude of difference between cohorts regardless of direction. Third, residualised violin plots were generated to visually inspect cohort differences in distributional shape after removing the effects of age and sex. For selected features, residuals were obtained by regressing the feature on age and sex separately within each cohort using a linear model, and the resulting residual distributions were compared between UKB and LIFE.

### 2.4 Hierarchical GAMLSS harmonisation

To harmonise neuroimaging features across cohorts while preserving biologically meaningful variability, we propose a framework based on Generalised Additive Models for Location, Scale, and Shape (GAMLSS) (Rigby & Stasinopoulos, 2005) with cohort-specific random effects. Existing ComBat-based approaches typically correct only the first two distributional moments under Gaussian or negative binomial assumptions. In contrast, the proposed framework can be implemented with any parametric distribution that has a tractable quantile function. Cohort effects are estimated within the relevant distributional parameters and then removed through quantile mapping, yielding harmonised values that preserve individual rank information and associations with biological covariates. The same modelling framework also provides normative deviation scores, allowing harmonisation and normative inference to be performed jointly within a single model.

#### 2.4.1 Distribution selection and model specification

Let *y_ij_* denote the observed value for individual from cohort . When the distributional family of a feature is known or can be specified from prior knowledge, that family could be specified directly. Otherwise, for neuroimaging features with unknown or non-Gaussian conditional distributions, we applied a prespecified distribution-selection hierarchy to identify an appropriate family.

The procedure begins with the four-parameter Sinh-Arcsinh distribution (SHASH) (Jones & Pewsey, 2009),

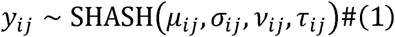

where *μij, σij, νij*, and τ*ij* represent the location, scale, skewness, and kurtosis parameters, respectively. SHASH was chosen as the primary candidate because it accommodates asymmetric and heavy-tailed distributions without requiring prior specification of the direction or degree of departure from normality, making it well suited to the diverse marginal distributions encountered across neuroimaging feature categories.

The location and scale submodels were specified consistently across all distributional families. Both include a cohort-specific random intercept, a nonlinear smooth effect of age estimated via penalised B-splines, and sex as a fixed effect,

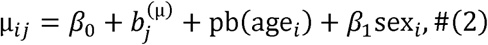

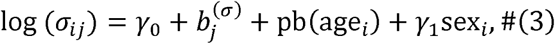

where 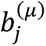 and 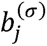 denote cohort-specific random intercepts, pb(·) denotes penalised B-spline smooth functions of age, and sex enters as a fixed effect in both parameters. The identity link function is applied to μ and the log link to σ to ensure positivity. For longitudinal applications, an additional subject-specific random intercept could be included in the location submodel to account for within-person dependence across repeated observations.

For the initial SHASH specification, the shape parameters were modelled with cohort-specific random intercepts to allow between-cohort differences in skewness and kurtosis beyond those captured by location and scale:

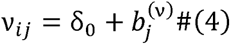

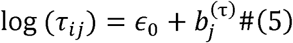

where 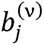 and 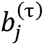 are cohort-specific random intercepts for skewness and kurtosis respectively.

If this specification fails to converge, simpler parameterisations are attempted sequentially by reducing one or both shape parameters to global intercept-only terms. If all SHASH specifications fail to converge, the framework falls back to the three-parameter Generalised Gamma distribution, which captures location, scale, and a single shape parameter without requiring a kurtosis term. If the Gamma distribution also fails, the Normal distribution is used as a final fallback. The distribution family assigned to each feature is reported in Supplementary Table 3. All models were fitted using the GAMLSS backfitting algorithm with a maximum of 200 iterations.

#### 2.4.2 Quantile mapping and cohort-effect removal

Harmonised values were obtained in three steps: first, cohort-specific centile scores were computed; second, harmonised distributional parameters were derived by removing cohort random effects; and third, quantile mapping was used to return values to the original measurement scale.

For each observation *y_ij_*, its centile score within its cohort-specific fitted distribution is computed using the cumulative distribution function (CDF) of the fitted family,

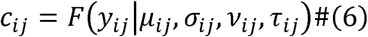

where the distributional parameters include all estimated cohort random effects. This centile score reflects the rank position of each observation within its own cohort’s distribution after adjusting for age and sex. It thus carries forward the individual-level biological information needed for subsequent quantile mapping.

Harmonised distributional parameters are then obtained by removing the cohort random effects from each distributional parameter. Because the random intercept *b_j_* are estimated under a zero-mean normal assumption, setting each random effect (*b_j_*) to zero on the link scale is equivalent to replacing each cohort’s distributional offset with the population-average effect. This is conceptually analogous to the empirical Bayes shrinkage employed in ComBat-based methods, where batch effect estimates are regularised toward a common prior mean rather than estimated independently per batch.

In both cases, the harmonisation operation replaces each cohort or site’s estimated distributional offset with the population-level expectation.

For a given distributional parameter *θ* with link function *g*(·), the corresponding harmonised parameter 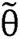 is defined as:

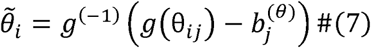

where 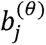 is the cohort-specific random intercept for that parameter. For the location parameter, this reduces to direct subtraction on the identity scale, whereas for scale and kurtosis the subtraction is performed on the log scale before exponentiation. Parameters for which no cohort random effect was estimated were carried forward without modification.

Harmonised values are then generated by applying the quantile function (inverse CDF) of the fitted distribution to each observation’s centile score under the harmonised parameters,

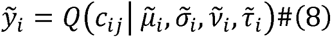

This operation preserves the rank ordering of each observation within its cohort while simultaneously removing cohort-specific distributional offsets from all fitted parameters.

Notably, because each centile score is obtained directly from the fitted conditional distribution, it can also be converted immediately into a normative deviation score without requiring a separate normative modelling step. Specifically, each centile score *c_ij_* is mapped to the standard normal scale via the probit transformation,

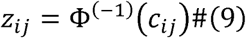

where 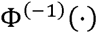 is the standard normal quantile function. This centile-to-z mapping is the conventional derivation of deviation scores in distributional normative modelling, including approaches based on fitted conditional distributions such as SHASH and related normative frameworks, where an individual’s covariate-adjusted percentile is converted to a z-score through the probability integral transform (de Boer et al., 2024; Dinga et al., 2021). Under a correctly specified model applied to a representative reference sample, these z-scores should follow a standard normal distribution.

#### 2.4.3 Post-processing of harmonised outputs

Following harmonisation, a structured quality control procedure was applied to each feature. First, observations with CDF values outside the open interval (0, 1) had their harmonised values set to missing, as values within this range are a necessary condition for valid quantile mapping. Second, harmonised values that were negative or infinite after back-transformation were also set to missing, as these outcomes are inconsistent with the non-negative measurement scale of the neuroimaging features analysed here. Finally, normative z-scores that were infinite or undefined following the probit transformation were set to missing.

### 2.5 Comparison models

To benchmark the proposed hierarchical GAMLSS harmonisation framework against established approaches, we compared its performance with three widely used ComBat-based methods: classical ComBat, ComBat-GAM, and ComBat-LS. These methods were selected because they represent the main progression of ComBat methodology from linear mean–variance adjustment, to nonlinear covariate modelling, to explicit location-scale correction. In contrast to the proposed framework, all three methods operate under Gaussian assumptions.

Classical ComBat modelled each feature using linear covariate effects with cohort-specific location and scale adjustment. ComBat-GAM extended this framework by modelling age nonlinearly using penalised B-splines, whilst retaining the same empirical Bayes site correction procedure. ComBat-LS further modelled site effects on both the location and scale parameters, with age again specified using penalised B-splines. All three ComBat variants were implemented using the neuroComBat R package (Fortin et al., 2017).

As the matrix computations in neuroComBat require complete datasets, we implemented a temporary imputation strategy for features with missing values. Missing data points were replaced with the per-feature mean prior to harmonisation. Following the computation of harmonised values, the original missing status was restored to ensure the analytic sample remained consistent across methods. We also removed any negative values from the ComBat-harmonised outputs to mirror the GAMLSS post-processing step.

### 2.6 Validation framework

To evaluate the performance of the proposed hierarchical GAMLSS harmonisation framework, we applied a two-component validation strategy assessing batch effect removal and biological effect preservation. All evaluations were performed on the aligned sample of participants with complete data across all harmonisation approaches, allowing direct like-for-like comparisons. Three ComBat-based methods served as benchmarks: classical ComBat, ComBat-GAM, and ComBat-LS, implemented as described in Section 2.5.

#### 2.6.1 Data retention

Data retention was assessed by quantifying the proportion of valid records lost following harmonisation relative to the pre-harmonisation data. Following harmonisation, values that were negative were set to missing, as these outcomes are physically invalid for the non-negative neuroimaging features examined in this study (volumetric measures in mm³, cortical thickness in mm, and surface area in mm²). A record was therefore classified as lost if it was valid before harmonisation but became missing afterwards. Quantifying data retention is a necessary component of the validation framework because harmonisation methods that operate under Gaussian assumptions may produce invalid negative values for features whose true distributions are bounded at zero (e.g., white matter hypointensity volume). This represents a direct cost of harmonisation that is not captured by other batch effect metrics.

#### 2.6.2 Batch effect removal

Batch effect removal was assessed using two complementary metrics applied to both pre-and post-harmonisation data, enabling direct before-and-after comparisons.

The primary metric was the cohort R^2^ increment, computed using the same general framework as the motivating analysis described in Section 2.3. For each feature, two ordinary least squares models were fitted: a biological model including age and sex, and an extended model additionally including cohort as a categorical predictor. In this validation analysis, age was modelled using a degree-3 polynomial to provide a more flexible representation of the nonlinear lifespan trajectory than in the motivating analysis. The increment in R^2^ when cohort was added to the biological model was interpreted as the proportion of variance attributable to cohort membership independent of age and sex.

As a complementary distributional measure, residual between-cohort differences were also quantified using the mean pairwise Kolmogorov-Smirnov (KS) statistic. For each feature, age and sex effects were first regressed out using a degree-3 polynomial model, and KS statistics were then computed for all unique cohort pairs on the resulting residuals. The mean KS statistic across cohort pairs was taken as the summary value for that feature. By residualising for biological covariates before testing, this analysis isolates distributional differences attributable to cohort rather than to differences in age or sex composition across samples. Unlike the R^2^ framework, which quantifies residual cohort-related variance in a regression setting, the KS statistic is sensitive to broader differences in distributional shape, including spread, skewness, and kurtosis. Results from both metrics are presented jointly to provide complementary assessments of mean-level and full-distributional batch removal.

#### 2.6.3 Biological effect preservation

An effective harmonisation method should remove technical between-cohort variation without attenuating biologically meaningful signal. Because age and sex are modelled as fixed biological covariates within the GAMLSS framework and should be preserved by design, their attenuation following harmonisation would indicate over-correction. We therefore evaluated preservation of two well-established sources of structural neuroimaging variation: age-related differences in brain morphology and sex differences in brain morphology. Both effects are expected across the cohorts included in the present study and are not nuisance signals that harmonisation should remove, making them suitable reference signals for preservation assessment.

##### Age effect preservation

For each feature, Spearman’s rank correlation with age was computed before and after harmonisation for each approach. Spearman correlation was selected in preference to Pearson correlation because it is robust to the non-Gaussian marginal feature distributions and does not assume a strictly linear relationship. This makes it well suited to monotonic age-related effects across the broad age range represented in the sample.

To complement this univariate analysis, a multivariate age-prediction model was fitted separately to each dataset using gradient-boosted regression trees (XGBoost) (T. Chen & Guestrin, 2016) with five-fold cross-validation. Predictive performance was summarised using mean absolute error (MAE) and R^2^. This multivariate analysis provides a complementary test of whether harmonisation preserves the joint age-related covariance structure across features, rather than only the marginal association of each feature with age considered in isolation.

##### Sex effect preservation

For each feature, the magnitude of sex differences before and after harmonisation was quantified using Cohen’s *d*, computed with the pooled standard deviation. Welch two-sample *t*-tests were additionally used to assess statistical significance, with false discovery rate (FDR) correction applied across features within each harmonisation approach. Results were summarised using the mean and median absolute Cohen’s *d* across features, the proportion of features with a significant sex difference (*q* < 0.05), and the proportion of features for which the absolute value of Cohen’s *d* decreased after harmonisation, as an indicator of sex effect attenuation.

##### Quantitative trajectory visualisation

To complement these quantitative metrics, age trajectories for each neuroimaging feature were visualised before and after harmonisation using binned cohort means (2-year bins) with Locally Estimated Scatterplot Smoothing (LOESS) smoothers (Cleveland, 1979), stratified by sex. These plots provided a qualitative check that cohort-level divergence in age trends was reduced after harmonisation without distorting the overall trajectory shape.

## 3. Results

### 3.1 Feature distribution characteristics

Full distributional moments by feature and cohort are reported in Supplementary Table 4 (original data) and Supplementary Table 5 (post within-cohort ComBat-Long harmonised data). Both tables show highly consistent patterns, indicating that the overall distributional structure of the features is largely preserved after within-cohort harmonisation.

Across the 237 structural neuroimaging features, marginal distributions were predominantly right-skewed and leptokurtic, with considerable variation across feature categories. In the post-ComBat-Long data, the median absolute skewness was 0.32 and the median excess kurtosis was 0.47 across all feature-cohort pairs. Global brain volumetric measures (total brain volume, supratentorial volume) and subcortical grey matter structures (hippocampus, amygdala, caudate, putamen, pallidum, thalamus) were approximately symmetric and near-Gaussian, with median absolute skewness below 0.5 and excess kurtosis close to zero (Supplementary Table 5). Cortical features also showed low rates of non-Gaussianity. The proportions of feature-cohort pairs exceeding absolute skewness of 1 and excess kurtosis of 3 were 1.8% and 3.3% for cortical volume, 1.2% and 3.4% for cortical thickness, and 3.5% and 6.3% for cortical surface area. An exception was left parahippocampal surface area in MAS, which showed marked positive skewness and heavy tails (skewness 12.17; excess kurtosis 297.42 in Supplementary Table 5).

According to Supplementary Table 5, the most pronounced departures from normality were concentrated in ventricular volumes and white matter hypointensity volume. Lateral and inferior lateral ventricle volumes showed skewness ranging from 1.09 to 3.86 and excess kurtosis up to 41.0 across cohorts. Similarly, the third ventricle showed skewness from 0.86 to 3.04 and excess kurtosis up to 30.1, and white matter hypointensity volume showed skewness from 1.06 to 7.95 and excess kurtosis up to 141.5 (compared to skewness 0.89 to 10.03 and excess kurtosis up to 207.4 in the original data, Supplementary Table 4). Notably, corpus callosum subregions also showed substantial non-Gaussianity, though this was highly cohort-specific. In both the original and post-ComBat-Long data, the mid-anterior, mid-posterior, and central subregions were severely right-skewed and leptokurtic in LIFE (skewness 5.04 to 8.52; excess kurtosis 103.8 to 239.8). MAS showed a secondary but considerably milder elevation (skewness up to 2.68; excess kurtosis up to 16.6), while the remaining four cohorts showed skewness below 1.3 and excess kurtosis below 3 for the same features. The posterior and pericallosal subregions were near-Gaussian across all cohorts.

### 3.2 Between-cohort distributional differences

Table 3 presents the variance in each neuroimaging feature attributable to biological covariates (age and sex) and to cohort membership, stratified by feature category. Two patterns are evident. First, cohort-related variance was strongly heterogeneous across feature categories. Second, in the most affected categories the batch effect was large relative to the biological signal.

Cortical thickness features were the most severely affected, with a median cohort R^2^ of 0.105 compared with a median biological R^2^ of only 0.045. Thus, the batch effect was on average approximately twice as large as the age and sex signal in cortical thickness, with 52.9% of features exceeding R^2^ = 0.10. Cortical volumes and subcortical structures showed intermediate and more modest batch effects overall, although left and right pallidum volumes were notable exceptions, with cohort R^2^ values of 0.301 and 0.199, corresponding to approximately 2.6 times their respective biological R^2^ values. Cortical surface area features showed negligible cohort-related variance, with a median cohort R^2^ of 0.005.

**Table 3.** Cohort effect R-squared by neuroimaging feature category.

| Feature category | n | Median $R^2$ (age + sex) | Median $R^2$ (cohort) | % $R^2 > 0.05$ | % $R^2 > 0.10$ | Max $R^2$ (cohort) |
| --- | --- | --- | --- | --- | --- | --- |
| Cortical Thickness | 68 | 0.045 | 0.105 | 69.1% | 52.9% | 0.284 |
| Cortical Volume | 66 | 0.126 | 0.037 | 27.3% | 4.5% | 0.185 |
| Subcortical/Global Volume | 37 | 0.213 | 0.016 | 16.2% | 8.1% | 0.301 |
| Cortical Surface Area | 66 | 0.152 | 0.005 | 4.5% | 1.5% | 0.149 |
| All features | 237 | 0.109 | 0.026 | 31.2% | 18.1% | 0.301 |

Table 4 examines whether between-cohort differences extended beyond location to scale, skewness, and kurtosis. Scale differences were the most widespread, with 51.1% of all features showing an absolute log(SD) ratio greater than 0.1 between UKB and LIFE, indicating at least an approximate 10% difference in spread. This pattern was most pronounced in cortical thickness, where 72.1% of features exceeded this threshold. The largest individual scale differences were observed in caudal anterior cingulate cortical thickness, where UKB showed approximately 68% greater spread than LIFE, and in left thalamus volume, where LIFE showed approximately 66% greater spread than UKB.

Skewness and kurtosis differences were more selective but remained substantial in specific feature types. In the subcortical and global volume category, 24.3% of features showed an absolute skewness difference greater than 0.5 and 32.4% showed an absolute excess kurtosis difference greater than 1, the highest proportions across all categories. Corpus callosum subregions showed the most extreme between-cohort differences, with mid-anterior and mid-posterior corpus callosum displaying skewness differences of 11.0 and 9.4, respectively. Excluding corpus callosum subregions, white matter hypointensity volume showed the largest skewness difference (0.99) and the largest excess kurtosis difference (27.0), with UKB showing substantially heavier tails than LIFE for this feature.

**Table 4.** Between-cohort differences in distributional spread and shape by neuroimaging feature category.

| Feature category | n | Median |  |  | % Feature |  |  |
| --- | --- | --- | --- | --- | --- | --- | --- |
|  |  | \|log(SD) ratio\| | \|Δ skewness\| | \|Δ kurtosis\| | \|log(SD) ratio\| > 0.1 | \|Δ skewness\| > 0.5 | \|Δ kurtosis\| > 1 |
| Cortical Thickness | 68 | 0.152 | 0.218 | 0.237 | 72.1% | 10.3% | 10.3% |
| Cortical Volume | 66 | 0.114 | 0.071 | 0.189 | 56.1% | 1.5% | 4.5% |
| Subcortical/Global Volume | 37 | 0.099 | 0.224 | 0.448 | 48.6% | 24.3% | 32.4% |
| Cortical Surface Area | 66 | 0.071 | 0.141 | 0.285 | 25.8% | 4.5% | 16.7% |
| All features | 237 | 0.101 | 0.148 | 0.250 | 51.1% | 8.4% | 13.9% |

Residualised violin plots visually confirmed these distributional differences. As illustrated for right rostral middle frontal cortical thickness (smri_thick_cdk_rrmdfrh) in Fig. 1, the residual distributions after removing age and sex effects differed clearly between UKB and LIFE in spread, with UKB showing a broader distribution and more pronounced tails in both sexes. Taken together, these findings indicate that cross-cohort batch effects were not restricted to location, but extended to scale, skewness, and kurtosis, providing empirical justification for a harmonisation framework that models cohort effects in higher-order distributional moments rather than in the mean alone.

**Fig. 1.**
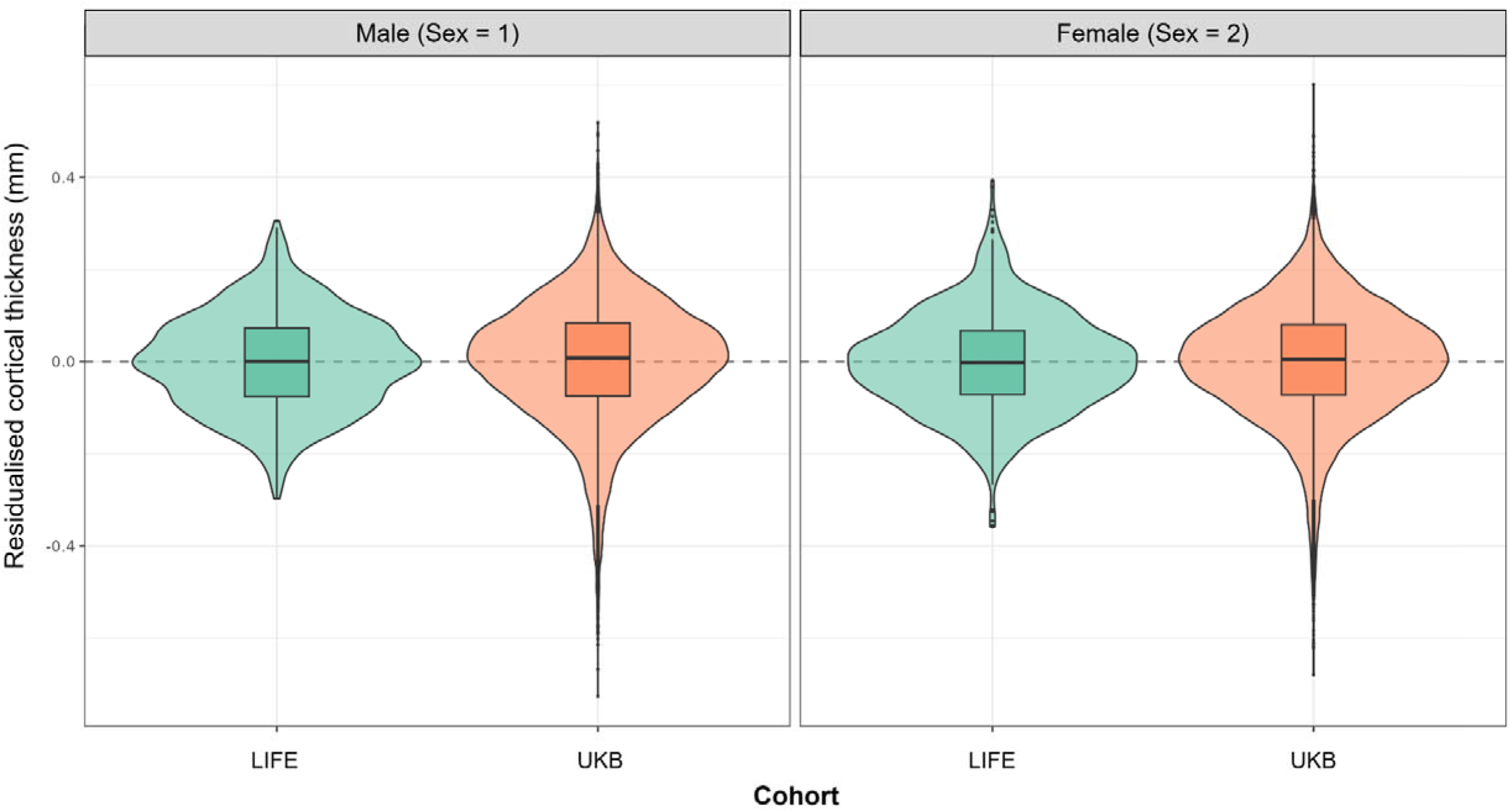
Residualised cohort distributions for right rostral middle frontal cortical thickness in UKB and LIFE. Violin plots show the distributions of residuals after adjustment for age and sex in baseline observations from UKB and LIFE restricted to their overlapping age range of 44 to 83 years. Residuals were obtained from cohort-specific linear models including age and sex and are shown separately for males (sex = 1) and females (sex = 2). Boxplots indicate the median and interquartile range. LIFE = Leipzig Research Centre for Civilization Diseases Adult Study; UKB = UK Biobank.

### 3.3 Data retention

The pre-harmonisation analytic dataset comprised 20,882,963 valid feature-level records across 88,126 observations and 237 neuroimaging features. The remaining 2,899 possible records, representing less than 0.01% of the full 88,126 × 237 matrix, were already missing before harmonisation because of the feature-level exclusions described in Section 2.2. Post-harmonisation data loss, defined as feature-level records that were valid before harmonisation but became missing afterwards because back-transformed values were negative or infinite, is summarised in Table 5.

**Table 5.** Post-harmonisation data retention pooled across all 237 neuroimaging features.

|  | ComBat | ComBat-GAM | ComBat-LS | GAMLSS |
| --- | --- | --- | --- | --- |
| Total valid records | 20,866,214 | 20,860,013 | 20,882,417 | 20,881,772 |
| Records lost, n | 16,749 | 22,950 | 546 | 1,191 |
| Records lost, % | 0.08% | 0.11% | <0.01% | 0.01% |
| Median NA rate across features | <0.01% | <0.01% | <0.01% | 0.01% |
| Max NA rate (single feature) | 16.2% | 13.4% | 0.3% | 0.4% |

Values are summarised across 20,882,963 feature-level records from 88,126 observations that were valid before harmonisation. Loss was defined as records that were valid pre-harmonisation but became missing after harmonisation because the corrected value was negative or infinite. Missing-value (NA) rates are reported across features, with the maximum value corresponding to the single feature with the highest post-harmonisation missingness. ComBat-GAM = ComBat with generalised additive modelling of age; ComBat-LS = ComBat with location-scale adjustment; GAMLSS = Generalised Additive Models for Location, Scale, and Shape.

Although data loss was negligible in absolute terms for all four methods, its magnitude and distribution differed substantially. ComBat-GAM produced the greatest total loss, with 22,950 records invalidated, corresponding to 0.110% of all pre-harmonisation valid records, across 193 of 237 features (81.4%). Classical ComBat showed a similar pattern, with 16,749 records lost (0.080%) across 187 features (78.9%). In both methods, losses were concentrated predominantly in the subcortical and global volume category. White matter hypointensity volume accounted for most of this loss, with per-feature invalidation rates of 16.2% under ComBat and 13.4% under ComBat-GAM. Inferior lateral ventricle volumes were also notably affected, with loss rates of 2.0% and 6.6% for the left side, and 0.9% and 3.6% for the right side, under ComBat and ComBat-GAM, respectively. By contrast, no feature in the cortical thickness, cortical volume, or cortical surface area categories exceeded a loss rate of 0.15% under either method. Per-feature missing-value rates for all methods are reported in Supplementary Table 8.

ComBat-LS produced the smallest total loss, with 546 records invalidated (0.003%) across 32 features (13.5%), and no individual feature showing a loss rate above 0.29%. GAMLSS invalidated 1,191 records (0.006%) across 200 features (84.4%), representing an approximately 14-fold reduction in total loss relative to ComBat and a 19-fold reduction relative to ComBat-GAM, although around twice the loss observed under ComBat-LS. Importantly, under GAMLSS no individual feature exceeded a loss rate of 0.43%, and no feature in any category crossed the 5% threshold. The broader spread of these minimal losses across features likely reflects the quantile-mapping procedure, in which a small number of back-transformed values fall outside the valid range at the extremes of the feature distribution. Unlike the ComBat-based approaches, where losses were concentrated in a small number of severely affected features, GAMLSS produced very small losses distributed more diffusely across features.

### 3.4 Batch effect removal

Batch effect removal was assessed using two complementary metrics: the cohort R^2^ increment (residual variance attributable to cohort after adjusting for age and sex) and the mean pairwise Kolmogorov-Smirnov (KS) statistic (residual cross-cohort distributional divergence after age and sex residualisation). Results are summarised in Table 6. Across both metrics, all four harmonisation methods reduced residual cohort effects relative to the pre-harmonisation data, but their effectiveness differed substantially.

#### Cohort R^2^ increment

Pre-harmonisation, cortical thickness features showed the largest batch effects, with a median cohort R^2^ increment of 0.097. Cortical volume, subcortical/global volume, and cortical surface area showed progressively smaller pre-harmonisation batch effects, with median R^2^ increments of 0.029, 0.030, and 0.010, respectively. Across all 237 features, the pre-harmonisation median cohort R^2^ increment was 0.030.

Following harmonisation, ComBat-GAM, ComBat-LS, and GAMLSS each achieved near-complete batch removal on this metric, reducing median cohort R^2^ increments to ≤ 0.001 across all feature categories. By contrast, classical ComBat achieved less complete correction, with median post-harmonisation R^2^ increments of 0.005 to 0.014 across categories. Feature-level results are reported in Supplementary Table 9.

#### Mean pairwise KS statistic

Before harmonisation, the median mean pairwise KS statistic across all features was 0.174, ranging from 0.106 in cortical surface area to 0.297 in cortical thickness. Thus, cortical thickness again showed the strongest residual cross-cohort divergence, with subcortical/global volumes also notably affected.

All methods reduced residual distributional divergence, but their effectiveness differed more clearly on this metric. ComBat-GAM achieved the lowest post-harmonisation KS values overall, with median values of 0.031 to 0.046 across feature categories. GAMLSS was the second most effective method, reducing the overall median KS statistic to 0.051 and clearly outperforming classical ComBat (median 0.122). ComBat-LS performed similarly overall (median 0.052), though it was weaker than GAMLSS for cortical thickness (0.068 vs 0.054) and subcortical/global volume (0.051 vs 0.047), while slightly stronger for cortical volume (0.050 vs 0.067). Per-feature KS results are reported in Supplementary Table 10.

**Table 6.** Residual batch effects before and after harmonisation, summarised by feature category.

| Feature category | n | Median cohort $R^2$ increment | | | | | Median mean pairwise KS statistic | | | | |
| --- | --- | --- | --- | --- | --- | --- | --- | --- | --- | --- | --- |
|  |  | Pre | ComBat | ComBat-GAM | ComBat-LS | GAMLSS | Pre | ComBat | ComBat-GAM | ComBat-LS | GAMLSS |
| Cortical Thickness | 68 | 0.097 | 0.014 | <0.001 | 0.001 | 0.001 | 0.297 | 0.149 | 0.041 | 0.068 | 0.054 |
| Cortical Volume | 66 | 0.029 | 0.005 | <0.001 | <0.001 | <0.001 | 0.162 | 0.108 | 0.031 | 0.050 | 0.067 |
| Subcortical/Global Volume | 37 | 0.030 | 0.012 | <0.001 | <0.001 | <0.001 | 0.210 | 0.146 | 0.046 | 0.051 | 0.047 |
| Cortical Surface Area | 66 | 0.010 | 0.005 | <0.001 | <0.001 | <0.001 | 0.106 | 0.090 | 0.034 | 0.043 | 0.042 |
| **All features** | **237** | **0.030** | **0.009** | **<0.001** | **<0.001** | **<0.001** | **0.174** | **0.122** | **0.036** | **0.052** | **0.051** |

Values are median cohort $R^2$ increments and median mean pairwise Kolmogorov-Smirnov (KS) statistics across features within each category. The cohort $R^2$ increment was defined as the increase in $R^2$ when cohort was added to an ordinary least squares model already containing age and sex. KS statistics were computed from age- and sex-residualised data for all unique cohort pairs and averaged within feature before summarisation by category. Lower values indicate more complete removal of residual cohort effects. Pre = Pre-harmonisation; ComBat-GAM = ComBat with generalised additive modelling of age; ComBat-LS = ComBat with location-scale adjustment; GAMLSS = Generalised Additive Models for Location, Scale, and Shape.

### 3.5 Biological effect preservation

Biological effect preservation was evaluated across three complementary measures: the univariate Spearman correlation between each feature and age, a multivariate age prediction model, and Cohen’s d for sex differences.

#### Age effect preservation

Supplementary Table 11 presents the per-feature Spearman rank correlations between individual features and age, computed before and after harmonisation for each approach. Across all 237 features, post-harmonisation age correlations were stronger than pre-harmonisation values under all four methods. GAMLSS achieved the highest mean and median absolute Spearman P across features (mean |P| = 0.473; median |P| = 0.516), followed closely by ComBat-LS (mean = 0.472; median = 0.512) and ComBat-GAM (mean = 0.469; median = 0.502). All four methods retained significant age associations (FDR-corrected q < 0.05) in 99.6%, with GAMLSS the only approach to achieve significance across all 237 features.

#### Multivariate age prediction

To evaluate whether harmonisation preserved the joint age-related covariance structure across features, an XGBoost age prediction model was trained and evaluated using five-fold cross-validation separately on each harmonised dataset. Results are summarised in Figure 2. GAMLSS achieved the lowest mean absolute error (MAE = 4.20 years; R^2^ = 0.945), followed by ComBat-LS (MAE = 4.22 years; R^2^ = 0.944), ComBat-GAM (MAE = 4.28 years; R^2^ = 0.929), and classical ComBat (MAE = 4.48 years; R^2^ = 0.929), compared with a pre-harmonisation MAE of 4.99 years (R^2^ = 0.930). All four methods improved on pre-harmonisation performance. Notably, ComBat-GAM and classical ComBat produced near-identical R^2^ values despite a 0.20-year difference in MAE, suggesting that ComBat-GAM improved local prediction accuracy without a corresponding gain in overall variance explained.

**Fig. 2.**
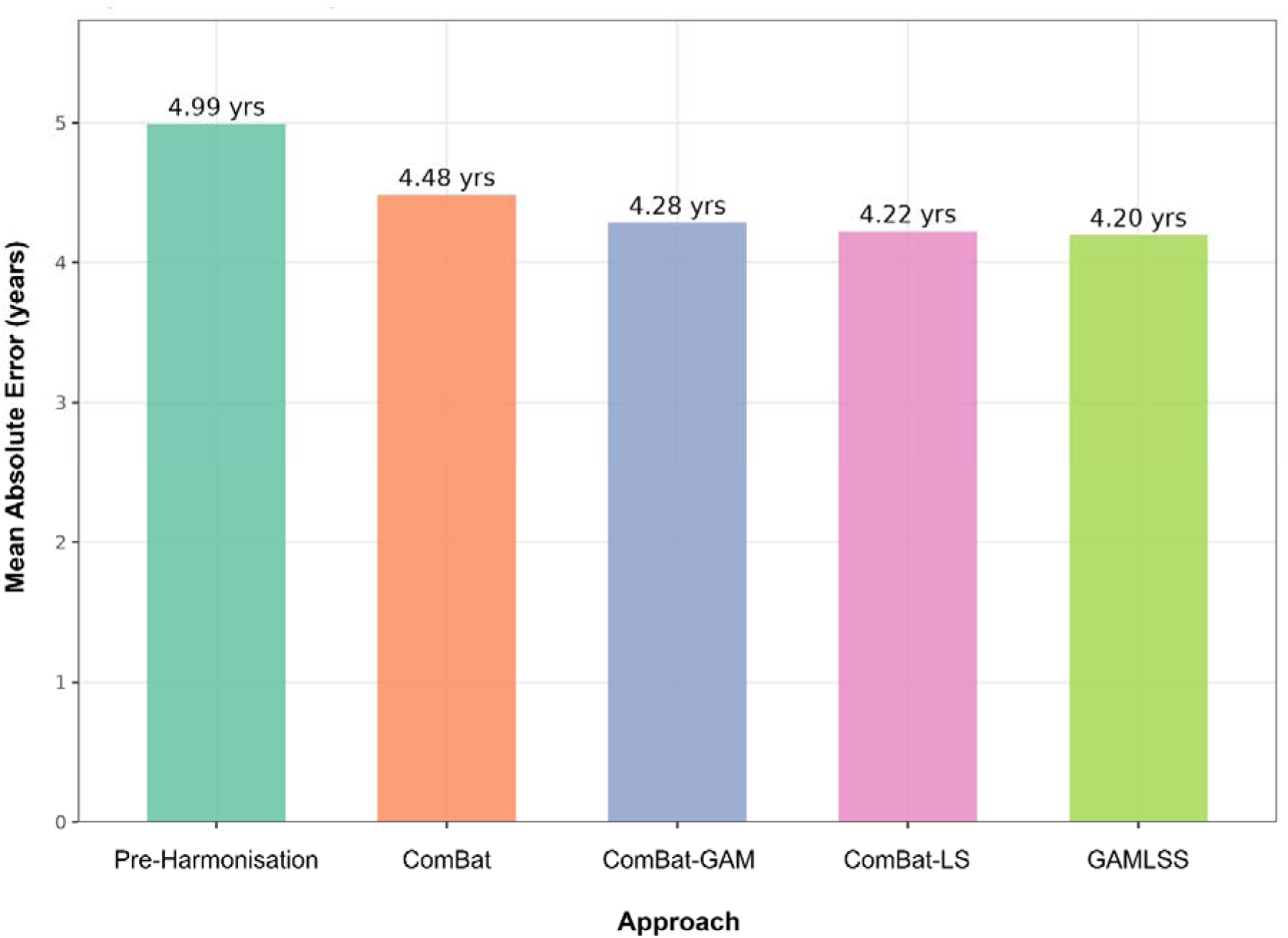
Multivariate age prediction performance by harmonisation method. Mean absolute error (MAE) from an XGBoost regression model trained on all 237 neuroimaging features jointly, evaluated using five-fold cross-validation. Lower MAE indicates greater retention of age-related biological information. Values above each bar indicate MAE in years. ComBat-GAM = ComBat with generalised additive modelling of age; ComBat-LS = ComBat with location-scale adjustment; GAMLSS = Generalised Additive Models for Location, Scale, and Shape.

#### Sex effect preservation

Cohen’s for sex differences was broadly preserved across all harmonisation approaches. Pre-harmonisation, the mean absolute Cohen’s across features was 0.450. Following harmonisation, mean absolute Cohen’s remained stable at 0.443 for ComBat, 0.434 for ComBat-GAM, 0.437 for ComBat-LS, and 0.437 for GAMLSS, corresponding to retention of 97-99% of the pre-harmonisation sex effect magnitude. The proportion of features with a statistically significant sex difference (FDR-corrected q < 0.05) was 99.2% before harmonisation and remained above 95.8% for all methods after harmonisation. No method showed systematic sex effect attenuation, and no feature showed an absolute change in Cohen’s d exceeding 0.20 under GAMLSS. Per-feature Cohen’s d values are reported in Supplementary Table 12.

#### Quantitative trajectory visualisation

To complement the quantitative metrics, age trajectories were visualised before and after harmonisation using binned cohort means (2-year bins) with LOESS curves, stratified by sex. Figures 3 and 4 illustrate trajectories for whole brain volume and white matter hypointensity volume respectively, chosen to represent a well-behaved near-Gaussian feature and the most severely non-Gaussian feature in the dataset.

For whole-brain volume (Fig. 3), pre-harmonisation trajectories showed clear cohort-level separation across the lifespan, with distinct offsets between cohorts in overlapping age ranges. Following harmonisation, ComBat reduced but did not fully resolve this separation, with UKB remaining visibly lower than LIFE and MAS across the overlapping adult age range in both sexes. ComBat-GAM, ComBat-LS, and GAMLSS produced substantially tighter cohort alignment, with broadly similar age trajectories across cohorts and preservation of the expected age-related decline in both sexes.

For white matter hypointensity volume (Fig. 4), pre-harmonisation trajectories showed broad cohort-level separation, particularly at older ages, together with the characteristic exponential increase with age. Following harmonisation, ComBat showed the clearest residual divergence between cohorts in overlapping age ranges, most notably between LIFE and UKB in adulthood. ComBat and ComBat-GAM both produced distorted trajectories in the younger cohorts, with IMAGEN and NCANDA showing erratic oscillations in the 10-to-25-year age range. In the ComBat-GAM panel, MAS additionally showed an altered trajectory shape, appearing more linear and steeply rising rather than exhibiting the exponential curvature seen in the pre-harmonisation data. Among the ComBat-based approaches, ComBat-LS preserved the most coherent overall distributional shape. However, the MAS trajectory appeared truncated at older ages. GAMLSS produced the most coherent trajectories overall, preserving the exponential increase across cohorts without visible distortion at younger or older ages.

**Fig. 3.**
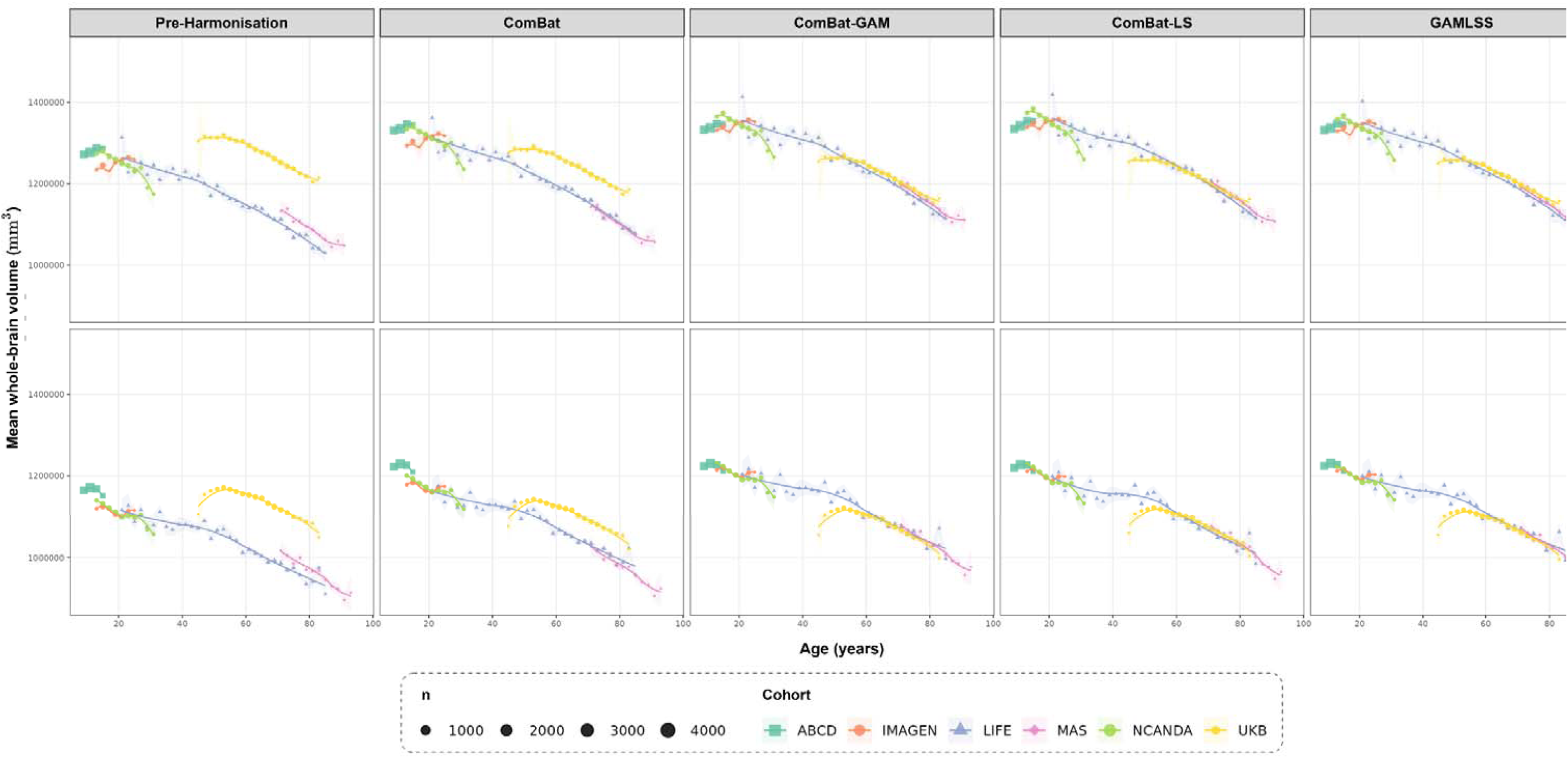
Age trajectories for whole-brain volume before and after harmonisation, stratified by sex. Each panel shows cohort-specific binned means (2-year bins) with standard error ribbons and LOESS curves. Point size reflects the number of observations per bin. Columns correspond to harmonisation method and rows correspond to sex. ComBat-GAM = ComBat with generalised additive modelling of age; ComBat-LS = ComBat with location-scale adjustment; GAMLSS = Generalised Additive Models for Location, Scale, and Shape.

**Fig. 4.**
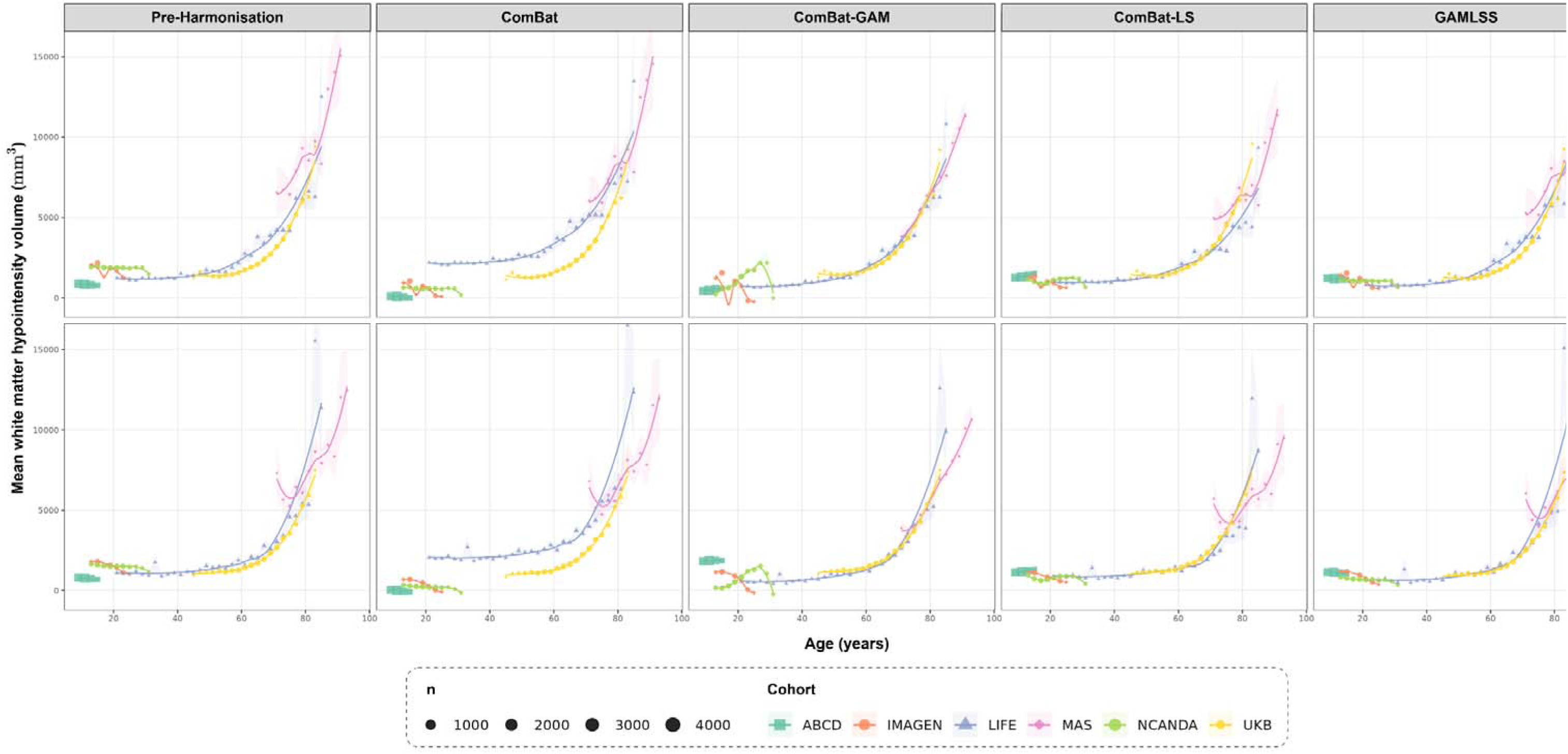
Age trajectories for white matter hypointensity volume before and after harmonisation, stratified by sex. Each panel shows cohort-specific binned means (2-year bins) with standard error ribbons and LOESS curves. Point size reflects the number of observations per bin. Columns correspond to harmonisation method and rows correspond to sex. ComBat-GAM = ComBat with generalised additive modelling of age; ComBat-LS = ComBat with location-scale adjustment; GAMLSS = Generalised Additive Models for Location, Scale, and Shape.

## 4. Discussion

In this study, we proposed a unified hierarchical GAMLSS framework for cross-cohort harmonisation and normative modelling of structural neuroimaging data and applied it to pooled MRI data from six cohorts spanning childhood to late life. By modelling cohort effects directly within the conditional distribution, the framework addresses a key limitation of existing approaches that either focus primarily on location-scale correction or produce deviation scores without harmonised outputs on the original scale. A central advantage of this framework is its distributional flexibility. Unlike ComBat and its variants, which operate under Gaussian assumptions, the proposed framework can be implemented with any parametric family for which exact inverse mapping is available. This makes it a more general tool for harmonisation of neuroimaging phenotypes whose conditional distributions may vary not only in mean and variance, but also in skewness and kurtosis. An additional strength is that normative deviation scores arise directly from the same fitted model, allowing harmonisation and normative inference to be conducted jointly rather than as separate downstream steps. This is particularly relevant for large-scale brain charting applications, where both harmonised native-scale measurements and covariate-adjusted deviation scores may be required. The empirical motivation analyses in Section 3.2 support the importance of this broader distributional view, showing that cross-cohort differences were not confined to location alone but extended to spread and, for selected feature classes, to higher-order moments.

The harmonisation performance of the proposed framework was evaluated across several complementary criteria, including data retention, residual batch effects, and preservation of biological signal. On data retention, GAMLSS and ComBat-LS each retained nearly all valid observations after harmonisation. This is consistent with the fact that both approaches accommodate features whose distributional shape departs from classical Gaussian correction. In contrast, the greatest data loss occurred under classical ComBat and ComBat-GAM, concentrated in strongly right-skewed and zero-bounded features such as white matter hypointensity and ventricular volumes. These findings suggest that data retention should be considered an integral component of harmonisation evaluation, as invalid values represent a direct analytic cost of imposing an inadequate distributional form on bounded neuroimaging outcomes.

Regarding batch-effect removal analysis, the proposed framework showed strong performance across both regression-based and distributional criteria. On the cohort R^2^ increment, GAMLSS reduced residual cohort-related variance to near-zero levels across all feature categories, performing comparably to ComBat-GAM and ComBat-LS and substantially better than classical ComBat. On the mean pairwise KS statistic, GAMLSS again showed marked improvement relative to the pre-harmonisation data and classical ComBat, with performance broadly similar to ComBat-LS. ComBat-GAM achieved the lowest post-harmonisation KS values overall, indicating the strongest residual cross-cohort alignment by that metric. However, this result should be interpreted alongside its poorer data-retention performance and the trajectory distortions observed for highly non-Gaussian features such as white matter hypointensity volume (Fig. 4). These findings suggest that ComBat-GAM may achieve stronger marginal distributional convergence partly by imposing a Gaussian location-scale correction on features whose support and shape depart substantially from that assumption. This improves KS performance, but at the cost of invalid values and reduced fidelity of the age trajectory for the most distributionally challenging phenotypes.

Biological signal preservation provided the clearest evidence for the advantages of the proposed framework. Across univariate age correlation, multivariate age prediction, and sex effect preservation, GAMLSS was consistently among the best-performing methods. The strengthening of age associations after harmonisation is interpretable as the unmasking of biological signal previously attenuated by cohort-level batch effects, consistent with the substantial residual cohort variance documented before harmonisation. This interpretation is further supported by the multivariate age prediction results, which test whether the joint feature-age covariance structure has been preserved rather than only marginal associations. The trajectory plots reinforced these findings. For a near-Gaussian feature such as whole-brain volume, GAMLSS produced cohort alignment comparable to the strongest comparators while preserving the expected age-related decline. For white matter hypointensity volume, a severely non-Gaussian feature, the advantages of a distribution-aware approach were more apparent: GAMLSS preserved the exponential age trajectory across cohorts without the visible distortions at younger or older ages seen under some ComBat-based approaches. This contrast is important, because it suggests that the main value of GAMLSS lies not simply in removing batch effects, but in doing so while better preserving biologically plausible trajectories for features whose conditional distributions are more difficult to model.

Several limitations should be acknowledged. First, although the study used longitudinal data from all six cohorts, the cross-cohort harmonisation performed here was effectively cross-sectional. The analysis was based on data that had already undergone within-cohort harmonisation using ComBat-Long before the cross-cohort GAMLSS model was fitted. This two-stage strategy was necessary because a single model that jointly accounted for within-cohort scanner effects, between-cohort differences, and subject-level repeated measures across all cohorts was not computationally feasible in the present setting. As a result, the current framework does not explicitly model within-subject dependence during the cross-cohort harmonisation step. This is an important limitation, but it is not unique to the present study. Fully integrated harmonisation across multiple waves, multiple scanners or sites within cohort, and multiple cohorts remains a major unresolved challenge in the field. The present framework therefore represents a pragmatic step rather than a complete solution to that problem.

Second, the computational cost of GAMLSS is substantially higher than that of the ComBat variants. Each feature requires a separate iterative model fit, and the burden increases with sample size, the number of parameters being modelled, and the complexity of the random-effects structure. This may limit feasibility in very high-dimensional settings or in applications where rapid harmonisation is required. Third, the current distribution-selection procedure remains heuristic. The sequential fallback from SHASH to Generalised Gamma to Gaussian provided a workable approach, but it is not yet a formally established model-selection strategy. More systematic approaches based on information criteria, residual diagnostics, or more efficient adaptive screening may improve both calibration and computational efficiency. Finally, the current model treats each neuroimaging feature independently and does not explicitly account for covariance across features. This means that while marginal distributions are harmonised, multivariate relationships may still be altered in ways that are not directly controlled.

These limitations point to several directions for future work. One major priority is the development of a truly joint longitudinal version of the framework that can accommodate subject-level repeated measures, scanner or site effects within cohorts, and cohort-level differences within a single model. Another important direction is improvement of the distribution-selection procedure, ideally through a more principled and computationally efficient strategy. Extending the framework beyond continuous outcomes will also be valuable. In particular, support for discrete, semi-continuous, or zero-inflated outcomes would broaden its applicability to a wider range of neuroimaging-derived variables. A further extension would be to develop multivariate variants that better preserve covariance structure across features, allowing harmonisation to target not only marginal distributions but also biologically meaningful inter-feature relationships. External validation in additional datasets and application to other imaging modalities will also help clarify the settings in which this framework provides the greatest advantage.

## 5. Conclusion

This study presented a unified hierarchical GAMLSS framework for cross-cohort harmonisation and normative modelling of structural neuroimaging data across six cohorts spanning the human lifespan. By modelling cohort effects across the full conditional distribution and returning harmonised values on the original measurement scale, the framework provides a flexible alternative to existing ComBat-based approaches, particularly for features with complex non-Gaussian distributions. Across multiple validation criteria, GAMLSS showed the strongest and most consistent preservation of biological signal, while also achieving substantial batch-effect removal and maintaining coherent post-harmonisation trajectories. Its ability to generate harmonised native-scale values and normative deviation scores within the same model further strengthens its practical value for large-scale neuroimaging studies. As multi-cohort population imaging studies continue to expand in size and complexity, flexible distribution-aware harmonisation frameworks of this kind may become increasingly important for ensuring that pooled analyses reflect biological variation rather than technical artefact.

## 6. Author contribution

M.P.H.: Conceptualisation; primary analysis; Writing – original draft. N.K.H., L.F.: Data analysis; Writing – review & editing. R.V., H.B., E.K.D., L.M.S.: Data preparation. J.J., P.S.S.: Supervision. W.W., L.M.: Conceptualisation; Supervision; Writing – review & editing.

## Acknowledgements

Data used in the preparation of this article were obtained from the Adolescent Brain Cognitive DevelopmentSM (ABCD) Study (https://abcdstudy.org), held in the NIMH Data Archive (NDA). This is a multisite, longitudinal study designed to recruit more than 10,000 children age 9-10 and follow them over 10 years into early adulthood. The ABCD Study® is supported by the National Institutes of Health and additional federal partners under award numbers U01DA041048, U01DA050989, U01DA051016, U01DA041022, U01DA051018, U01DA051037, U01DA050987, U01DA041174, U01DA041106, U01DA041117, U01DA041028, U01DA041134, U01DA050988, U01DA051039, U01DA041156, U01DA041025, U01DA041120, U01DA051038, U01DA041148, U01DA041093, U01DA041089, U24DA041123, U24DA041147. A full list of supporters is available at https://abcdstudy.org/federal-partners.html. A listing of participating sites and a complete listing of the study investigators can be found at https://abcdstudy.org/consortium_members/. ABCD consortium investigators designed and implemented the study and/or provided data but did not necessarily participate in the analysis or writing of this report. This manuscript reflects the views of the authors and may not reflect the opinions or views of the NIH or ABCD consortium investigators. This work was supported by funding from the NIH (R01AA030575, Mewton/Squeglia; and K24AA031052, Squeglia).

This research was also conducted using data from the UK Biobank (Application Number 98013), and computational resources provided by the National Computational Infrastructure (NCI Australia), and the Katana cluster, which is supported by Research Technology Services at UNSW Sydney (Katana, 2010). M.P.H. is supported by the University International Postgraduate Award, University of New South Wales, Sydney. We acknowledge the use of Claude AI for assistance with grammar and language editing. All scientific content, analyses, and interpretations remain the authors’ original work.

## 7. Code availability

The framework for fitting and generating harmonised normative deviation score results is publicly available on GitHub at: https://github.com/maiho24/gamlssHarmo

